# Post-treatment recovery of docetaxel-treated prostate cancer monolayer and spheroid culture

**DOI:** 10.1101/2025.11.17.688723

**Authors:** Steph Swanson, Catherine B. McKenna, Elahe Cheraghi, Ernest Osei

## Abstract

In the last half-century, it has become widely recognized that small 3D aggregates of cancer cells called tumor spheroids mimic some aspects of tumor behavior. Cell culture geometry has been shown to influence drug pharmacokinetics, delivery, and resistance. Despite the improved physiological relevance, the laborious nature of spheroids has limited clonogenic measurement and longitudinal observation of post-treatment recovery in docetaxel-treated prostate tumor spheroids. Agent-based modeling can complement spheroid experiments by probing questions of interest that are experimentally inaccessible. Here, we performed proliferation and clonogenic assays in docetaxel-treated PC3 cells cultured in monolayers and spheroids to assess and compare end-of-treatment survival and post-treatment recovery. We observed growth stimulation with no survival benefit in low dose docetaxel-treated monolayer and spheroid culture. However, agent-based modeling suggested that this hormetic effect may have been influenced by the active process of apoptosis. To the best of our knowledge, this is the first clonogenic measurement of docetaxel-treated spheroid culture and longitudinal observation of post-treatment docetaxel-dose dependent effects in prostate cancer cell culture.

## 1 Introduction

The most prevalent cancer among North American men [1, 2], prostate cancer is clinically heterogeneous. Patients generally present with an indolent and localized disease that has a high cure rate and 5-year survival [3], but a subset of patients exhibit aggressive disease with progression and metastasis [4]. The standard of care for patients with recurrent, locally advanced, or metastatic prostate cancer is medical castration via androgen deprivation therapy (ADT) [5]. However, ADT fails in a median of 18–24 months and patients inevitably develop castration resistant prostate cancer [6]. With the publication of two randomized phase 3 trials in 2004 [7, 8], docetaxel (DTX) chemotherapy emerged as a new standard of care for metastatic castration resistant prostate cancer patients. Subsequently, DTX treatment was additionally investigated in hormone sensitive prostate cancer patients and found to benefit those with high-volume metastatic disease [9–11]. As such, DTX is the first-line chemotherapeutic drug in advanced prostate cancer [5].

Following the observation of paclitaxel cytotoxicity in the late 1960s [12], DTX was discovered in 1986 via extraction from the *Taxus baccata* yew tree [13] as another member of the chemotherapeutic taxane family. Both taxanes stabilize microtubules, which prevents depolymerization during mitosis and causes cell cycle arrest in the G2/M phase via the mitotic spindle-assembly checkpoint [14, 15]. Senescence can arise from permanent cell cycle arrest [16], but arrest can also be transient if the checkpoint releases without the necessary repair [17]. In this case, cells can experience mitotic slippage and enter a tetraploid or “pseudo-G1” phase without physically splitting into two daughter cells, leading to eventual apoptosis [15]. If cell division occurs with improperly segregated chromosomes, aberrant mitosis leads to mitotic catastrophe [18]. These non-viable cells can undergo a necrosis-like cell death after developing micronuclei as nuclear envelopes surround individual or groups of chromosomes [16]. That said, apoptosis has also been observed following mitotic catastrophe [19].

The mechanism of DTX-induced cell death has been largely observed through conventional 2D monolayer cell culture. However, cell-cell and cell-environment interactions in small 3D aggregates of cancer cells called tumor spheroids are more physiologically representative of patient tumors compared to monolayer architectures [20, 21]. Unlike the homogeneous nature of monolayer culture, heterogeneous cell populations arise in spheroids due to diffusion-limited nutrient and waste distributions [22]. These factors influence cellular morphology [23], signaling [24], proliferation [25], gene and protein expression [26], as well as drug pharmacokinetics [27], delivery [28], and resistance [29]. Consequently, there has been significant interest in using spheroid culture as an *in vitro* platform for drug discovery, testing and characterization [23, 27, 28, 30–32].

Nevertheless, the use of tumor spheroids for drug investigation is known to be tedious and time consuming [30]. Drug testing in spheroid cultures also suffers from sensitivity to spheroid size [32] and lack of reproducibility [31]. As such, thorough investigation requires analysis of different drug dosages, spheroid sizes, drug penetration into the spheroid, and impact of the spheroid environment on drug delivery. Moreover, some questions of interest cannot be accessed experimentally with adequate accuracy, particularly those related to the behavior of individual cells within the spheroid structure. This laborious combinatorial experiment design and experimental inaccessibility motivates the supplemental use of well informed *in silico* models [33]. Drug efficacy is influenced by mechanical and molecular tumor spheroid characteristics, which have been extensively modeled mathematically with *in silico* methods [34]. Agent-based modeling (ABM), a discrete *in silico* method, simulates these characteristics by considering each cancer cell as an individual “agent” that responds to its local environment, including neighboring cells, nutrients, and drugs [35, 36]. The rule-based structure of ABM is well suited for performing logical experiments with perfectly understood biological mechanisms. Under the assumption of accurate parameter estimates informed by *in vitro* and/or *in vivo* measurements, *in silico* models have particular utility in mechanistic hypothesis testing and falsification [37]. With respect to chemotherapy investigation, *in silico* methods also reduce exposure to dangerous chemicals.

There has been substantial work elucidating the mechanism of DTX [14, 38, 39]. More recent work has thoroughly characterized end-of-treatment survival in prostate cancer monolayers by proliferation assay [40–42], and to a lesser extent, clonogenic assay [43–45]. DTX-treated prostate cancer spheroids have also been investigated via proliferation assay [46, 47]. However, post-treatment recovery in the week following DTX exposure has received little investigation, despite the sometimes delayed outcomes associated with DTX-induced cell cycle arrest. Proliferation assays like the widely used, non-toxic Alamar Blue (AB) measure fluorescence produced by enzyme activity associated with cellular metabolism [48]. Often employed directly at end-of-treatment, proliferation assays can also monitor longitudinal changes in cellular activity. In contrast, clonogenic assay plates cells at low density to identify those capable of colony formation via continuous reproduction and thus, directly measures clonogenic potential [49]. Clonogenic assay is typically employed to quantitatively estimate survival fraction by colony counting, but post-treatment behavior can also be qualitatively investigated through longitudinal observation of colony formation. Here, we assessed end-of-treatment survival and post-treatment recovery of DTX-treated PC3 prostate cancer cells cultured in monolayers and spheroids to measure immediate and delayed effects of DTX treatment. Measurements were performed with clonogenic and AB proliferation assays to assess cell viability and activity, respectively. Qualitative and quantitative results were compared with fluorescence microscopy (FM) and ABM simulations.

## 2 Methods

### 2.1 Monolayer and spheroid cultures

PC3 cells (human prostate adenocarcinoma) were cultured in RPMI 1640 (Sigma-Aldrich, USA) culture medium supplemented with 10% fetal bovine serum (Thermo Fisher Scientific, USA) and 1% penicillin (Thermo Fisher Scientific, USA) and incubated in a humidified atmosphere at 37^*°*^C and 5% CO_2_. On day 0 of the experiment, monolayer cell culture was plated in T-25 culture flasks. Concurrently, prostate tumor spheroids were seeded and cultured in 2% agarose 3D Petri Dish (MicroTissues Inc., USA) gels. The 9 *×* 9 array of wells contained in each agarose gel was seeded with a suspension of PC3 cells such that spheroids of approximately 1,000 cells formed in each well (Fig. 1A).

**Figure 1.**
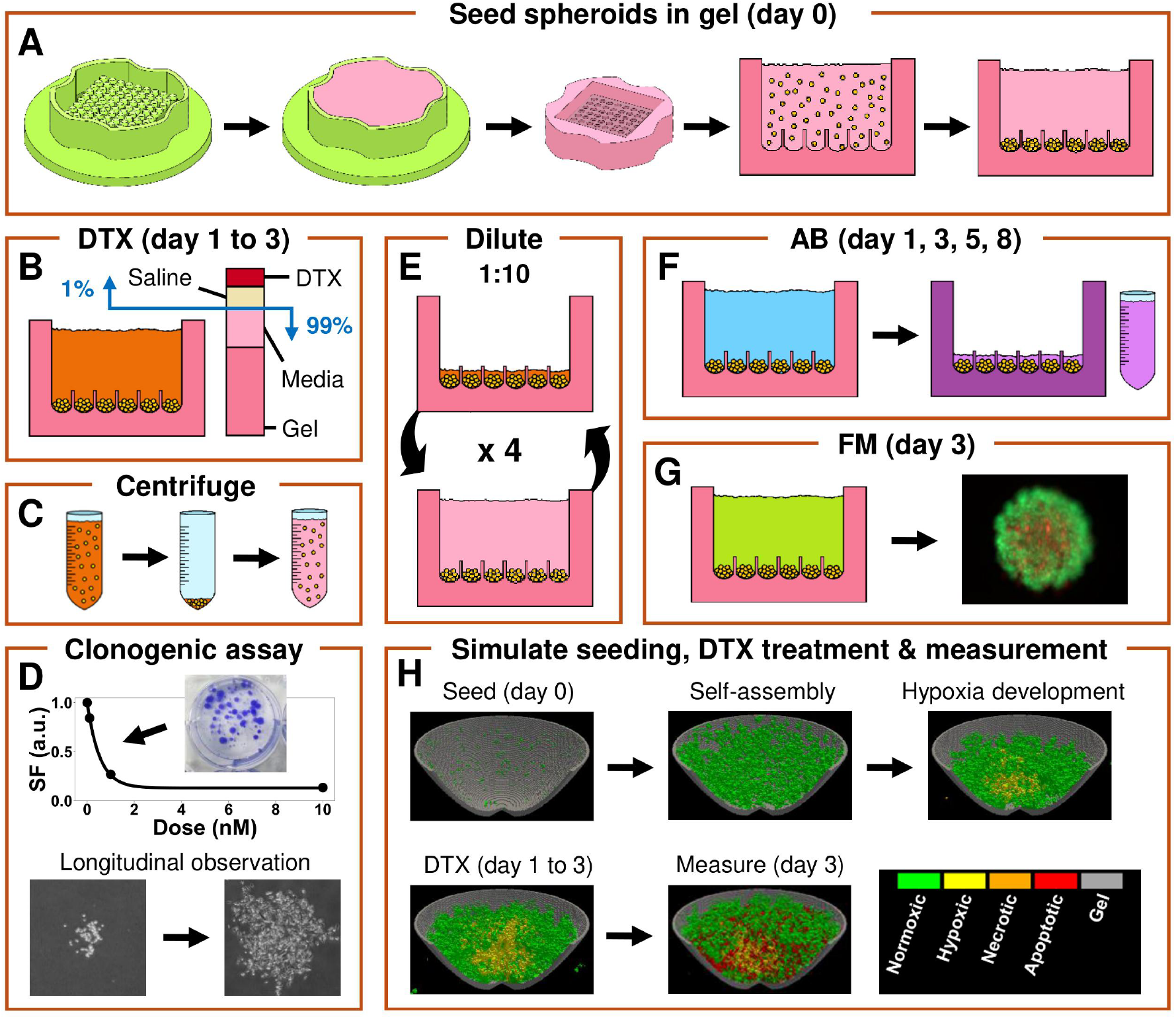
(A) Cells (yellow) seeded into agarose gels (pink) created from molds (green) self-assemble into prostate tumor spheroids in the bottom of wells. (B) Spheroids were exposed to 0, 0.1, 1, and 10 nM DTX diluted in saline with a 1:100 ratio of DTX in saline to culture medium and gel for 48 hours. DTX was removed by (C) centrifuging for (D) SF calculation and longitudinal observation via clonogenic assay, or by (E) serial dilution of spheroid-containing gels for (F) AB proliferation assay and (G) FM. (H) 3D ABM of spheroid seeding, DTX treatment, and measurement applied fitted clonogenic survival from experiment during treatment. DTX: docetaxel; SF: survival fraction; AB: Alamar Blue; FM: fluorescence microscopy; ABM: agent-based modeling.

### 2.2 Docetaxel chemotherapy treatment

24 hours post-seeding, monolayer and spheroid cultures were exposed to 0, 0.1, 1, and 10 nM DTX diluted in 0.9% NaCl in water (saline) for a duration of 48 hours (Fig. 1B). At each dose, a 1:100 ratio of DTX in saline to culture medium was maintained, with control samples exposed to 1:100 of saline to culture medium. In spheroid culture, the 500 *µ*L gel was included in the culture medium volume for calculating treatment concentration. After the 48 hour treatment period, measurement was performed. For monolayer culture, DTX treatment solution was removed from culture flasks which were washed with PBS (Wisent Inc., Canada) before replacement with supplemented culture medium. For spheroid culture, the DTX removal process depended on the measurement. For clonongenic assay, which requires spheroids to be disaggregated into a cell suspension, DTX was removed by centrifuge (Fig. 1C). However, AB proliferation assay and FM require intact spheroids to remain in the agarose gel, which retains its concentration of DTX after removal of surrounding treatment media. As such, DTX removal for these methods was achieved by serial dilution (Fig. 1E): all liquid was carefully removed from around and inside the gel and replaced drop-wise with 5 mL of RPMI, then gels were left to soak for one hour. The process was repeated four times so that the DTX in each 500 *µ*L gel was diluted by a factor of 10,000. In the final dilution before incubation, spheroid gels received supplemented culture medium.

### 2.3 Clonogenic assay

Clonogenic assay was performed with monolayer and spheroid cultures immediately after the 48 hour treatment period to measure clonogenic survival and observe longitudinal colony formation (Fig. 1D). Our detailed protocol for spheroid disaggregation is described elsewhere [50]. Briefly, for each dose of DTX, all spheroids were transferred from gels into a falcon tube for mechanical vortexing and incubation for two hours. Then, supernatant was removed and replaced with 0.25% trypsin-EDTA (Thermo Fisher Scientific, USA) before ten minutes of incubation, mechanical vortexing, and centrifuging for cell counting. Monolayer culture was incubated with 0.25% trypsin-EDTA for 3 minutes then centrifuged for cell counting. Counted cells from each dose were plated at low densities in 6-well plates for approximately two weeks of growth. Bright-field microscopy images of plated colony formation were taken regularly with an Olympus CKX53 inverted microscope. For colony counting, plates were fixed with 4% formaldehyde (Thermo Fisher Scientific, USA) in PBS for 1 hour then stained with 0.5% crystal violet (Thermo Fisher Scientific, USA) dissolved in water for 30 minutes. The plating efficiency (PE) was calculated using control samples as follows:

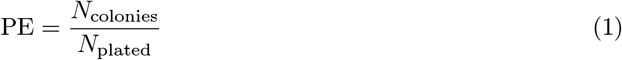

Here, *N*_colonies_ and *N*_plated_ are the number of colonies counted and plated, respectively. Survival fraction (SF) was calculated using PE as follows:

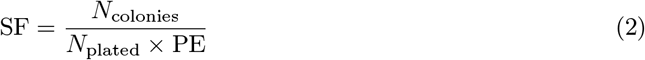

We developed the following novel model for least-squares fitting of SF values for each dose *d*:

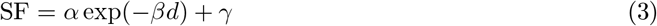

with parameters *α, β*, and *γ* extracted separately for monolayer and spheroid cultures, under the assumption that *α* + *γ* = 1. That is, the fitted curve is assumed to predict a SF of 1 at zero dose.

### 2.4 Alamar Blue proliferation assay

AB proliferation assay was performed with monolayer and spheroid cultures to measure average cellular proliferation pre-treatment on day 1, immediately after the 48 hour treatment period on day 3, and throughout post-treatment recovery on days 5 and 8 (Fig. 1F). Cells were not removed from their culturing vessels between treatment and AB measurement. Therefore, T-25 culture flasks for AB readings of monolayer culture were seeded according to dose (Table 1). Control samples were seeded as sparsely as possible to prevent the monolayer from becoming confluent by day 8, while ensuring a sufficient sample of cells for the pre-treatment measurement. Monolayers treated with the highest dose of 10 nM were seeded as densely as possible to ensure a sufficient number of cells remained adhered post-treatment, while still preventing confluency at the pre-treatment measurement. In contrast, all spheroids were formed of approximately 1,000 cells.

**Table 1.** Cells seeded AB proliferation assays of monolayer and spheroid cultures per T-25 flask and well in 3D Petri Dish gel, respectively.

| DTX dose (nM) | Cells/T-25 flask (monolayer) | Cells/well (spheroid) |
| --- | --- | --- |
| 0 (control) | 13,000 | 1,000 |
| 0.1 | 13,000 | 1,000 |
| 1 | 19,000 | 1,000 |
| 10 | 400,000 | 1,000 |

Prior to each AB measurement, samples were incubated in 5% AB (Sigma-Aldrich, USA) in RPMI for 3 hours at 37^*°*^C with 5% CO_2_ in a humidified atmosphere. For spheroids, culture medium was first removed from around and inside the gel, then 190 *µ*L of AB solution in RPMI was added to achieve the 5% concentration throughout the 500 *µ*L gel. Following incubation, AB solution was extracted and its absorbance intensity recorded at 570 nm using a BioTek Synergy H1 microplate reader. For each experimental condition, three 100 *µ*L aliquots of AB solution were removed from one monolayer-containing T-25 flasks and three spheroid-containing gels. Each AB measurement day, T-25 flasks and gels without cells were processed identically and measured to determine background absorbance, which was subtracted from that day’s monolayer and spheroid measurements, respectively. Due to the different seeding conditions, each experimental condition was then normalized to its day 1 measurement.

### 2.5 Fluorescence microscopy

FM was performed immediately after the 48 hour treatment period to visualize cell death within spheroids using the Live/Dead Cell Viability assay (Sigma-Aldrich, USA) (Fig. 1G). Live cells were labeled green with calcein acetoxymethyl (calcein-AM) [51] and dead cells were labeled red using propidium iodide (PI) [52]. Each gel was carefully emptied of culture medium and incubated for one hour with 0.2875 *µ*L calcein-AM, 1.15 *µ*L PI, 95 *µ*L RPMI, and 95 *µ*L PBS. Then, a Zeiss Axio Observer widefield microscope was used to capture fluorescent images of spheroids in gels that had been transferred to coverslip-bottomed Petri dishes. The microscope was operated by ZEN2 Blue Edition software and equipped with an Axiocam 506 mono camera and a Plan-Apochromat 63/1.4 Oil Ph3 M27 objective (Carl Zeiss Microscopy LLC, USA). An incubation plate (PECON, Germany) maintained the temperature at 37^*°*^C during imaging.

### 2.6 Simulation

In this study, an ABM of 3D tumor spheroid growth developed previously [50] was adapted to simulate DTX treatment (Fig. 1H). Briefly, 1,000 well oxygenated *Normoxic* tumor cells were placed randomly within a curved well inside the agarose gel. The cells fell to the bottom of the well under gravity and, as living cells energetically prefer contact with other living cells, self-assembled into a spheroid. A constant partial pressure of oxygen was maintained in the culture medium. Controlled through a partial differential equation (PDE), oxygen diffused through the spheroid and was consumed by cells. The local partial pressure of oxygen determined cell growth, with volume increasing according to a Michaelis-Menten saturation curve. Once a cell achieved a sufficient volume, it performed cell division. Oxygen-deprivation due to diffusion-limited oxygenation and competition caused *Normoxic* cells to reversibly become *Hypoxic*. Under severe oxygen deficiency, cells irreversibly became *Necrotic*. The 5 *µ*m-voxelized simulation space was 810 *µ*m across and 405 *µ*m deep, and the simulation time scale was determined such that *Normoxic* cells performed cell division approximately every 24 hours. The shape of the curved well and dimensions of simulated spheroids after 24 hours were calibrated to those in OCT images [53]. Then, 24 hours post-seeding, DTX treatment was simulated through a new PDE that was added to the model. Throughout the 48 hour treatment period, the simulated dose was maintained in the culture medium and gel and diffused through the spheroid with a diffusion coefficient of 256 *µ*m^2^ s^*−*1^ employed by Lai and Friedman [54] and Gordon [55]. During treatment, when cells attempted to perform cell division, they succeeded probabilistically according to the fitted curve (Eq. 3) derived from spheroid clonogenic experiments. Cells that failed to divide irreversibly became *Apoptotic*.

Following the 48 hour treatment period, numerical analysis of simulated spheroids treated with 0, 0.1, 1, and 10 nM DTX was performed. Simulated clonogenic SF was calculated using Eq. 2 by estimating *N*_colonies_ and *N*_plated_ as the number of living cells and the total number of cells in the spheroid, respectively. Simulated AB SF was calculated by estimating cellular activity of living and dying cells and normalizing to control. Living cells contributed to the calculation through the sum of each cell’s oxygenation. *Apoptotic* cells were assumed to be in cell cycle arrest for 24 hours following failed cell division [18], before undergoing the active process of apoptosis for 24 hours [56]. Cells actively undergoing apoptosis contributed to the AB estimation by assuming an equivalent level of activity as a fully oxygenated living cell. Simulated growth was estimated as the sum of each cell’s oxygenation normalized to that of control, and thus represents the simulated AB SF value without the contribution of active apoptosis.

Each dose was simulated ten times for statistical analysis, as the Monte Carlo simulation is stochastic in nature. A detailed mathematical description of the model can be found in the supplementary data.

### 2.7 Statistics

For comparison with clonogenic SF measurements, post-treatment day 3 AB absorbance was further normalized to that day’s control measurement. As AB measurements were performed in triplicate, individual values with obviously high leverage (i.e. that exerted undue influence on two similar values) were removed. All measurements are presented as averages with error bars. As described by Gupta et al. [57], clonogenic SF error was estimated by Fieller’s theorem using a 95% confidence interval from a Poisson distribution. All other error bars depict standard deviation. Clongenic EC_50_ values for monolayer and spheroid cultures were determined by identifying the dose that yielded a SF of 0.50 according to the model fit given by Eq. 3.

## 3 Results

Figure 2 summarizes end-of-treatment survival measurements performed immediately after the 48 hour DTX treatment period. Figure 2A shows that compared to monolayer culture, clonogenic survival of spheroid culture was comparable at 0.1 nM DTX, lower at 1 nM DTX, and higher at 10 nM DTX, with good fit to Eq. 3 for each culture type. Clonogenic EC_50_ was 0.76 nM and 0.45 nM DTX in monolayer and spheroid culture, respectively. Experimental and simulated clonogenic and AB spheroid measurements immediately post-treatment period are given in Figure 2B. Simulated spheroid clonogenic survival agrees well with experimental measurement and both significantly decrease with increasing dose. In contrast, we observed that with increasing DTX dose, experimental and simulated AB measurement increased before decreasing to minima of 87% and 89% of control at 10 nM, respectively (Fig. 2B). Simulated growth, which removes the contribution of apoptotic activity from the simulated AB calculation, is comparable to simulated clonogenic survival (Fig. 2B).

**Figure 2.**
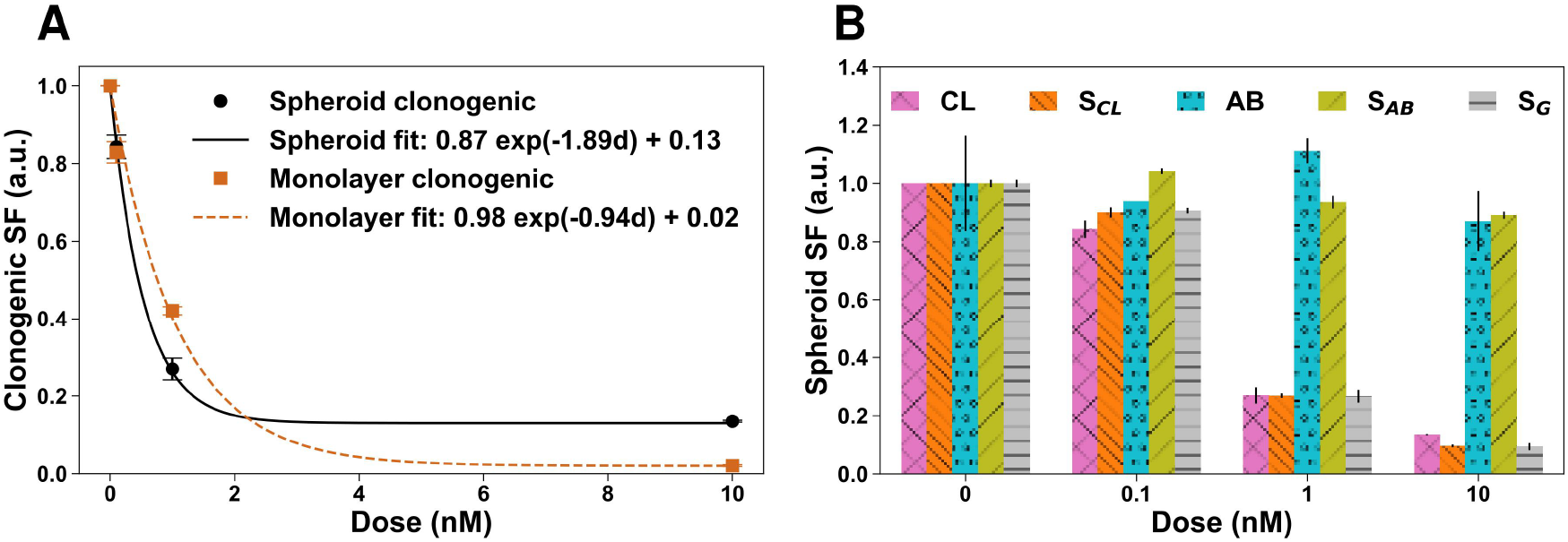
Measurements performed immediately after the 48 hour treatment period with 0, 0.1, 1, and 10 nM DTX. (A) Clonogenic assay SF values with model fit line given by Eq. 3 for 3D spheroids (black circle, solid line) and 2D monolayer culture (orange square, dashed line). (B) SF of spheroids calculated via clonogenic assay measurement (purple x hatch), simulated clonogenic assay S_*CL*_ (orange \\ hatch), normalized absorbance from AB proliferation assay measurement (blue circle hatch), simulated AB proliferation assay S_*AB*_ (olive / hatch), and simulated growth S_*G*_ (grey - hatch). Data shown are averages with error bars as described in Methods Section 2.7. DTX: docetaxel; SF: survival fraction; CL: clonogenic assay; AB: Alamar Blue; G: growth.

Figure 3 shows FM and simulated images of spheroids immediately post-treatment. In the first row, FM enface images of all DTX doses reveal peripheral green intensity with a faint central red spheroid core in control that increases with dose (Fig. 3A-D). The second and third rows of Figure 3 show simulated enface and volumetric images, respectively. The simulated control spheroid contains a yellow hypoxic core surrounding green proliferating *Normoxic* cells (Fig. 3E,I). In the simulated spheroid treated with 0.1 nM DTX, red *Apoptotic* cells appear sparsely throughout the hypoxic spheroid bulk (Fig. 3F,J). At higher doses of 1 nM and 10 nM DTX, there is no hypoxia and the majority of the spheroid is comprised of red *Apoptotic* cells with few green *Normoxic* cells throughout (Fig. 3G-H,K-L).

**Figure 3.**
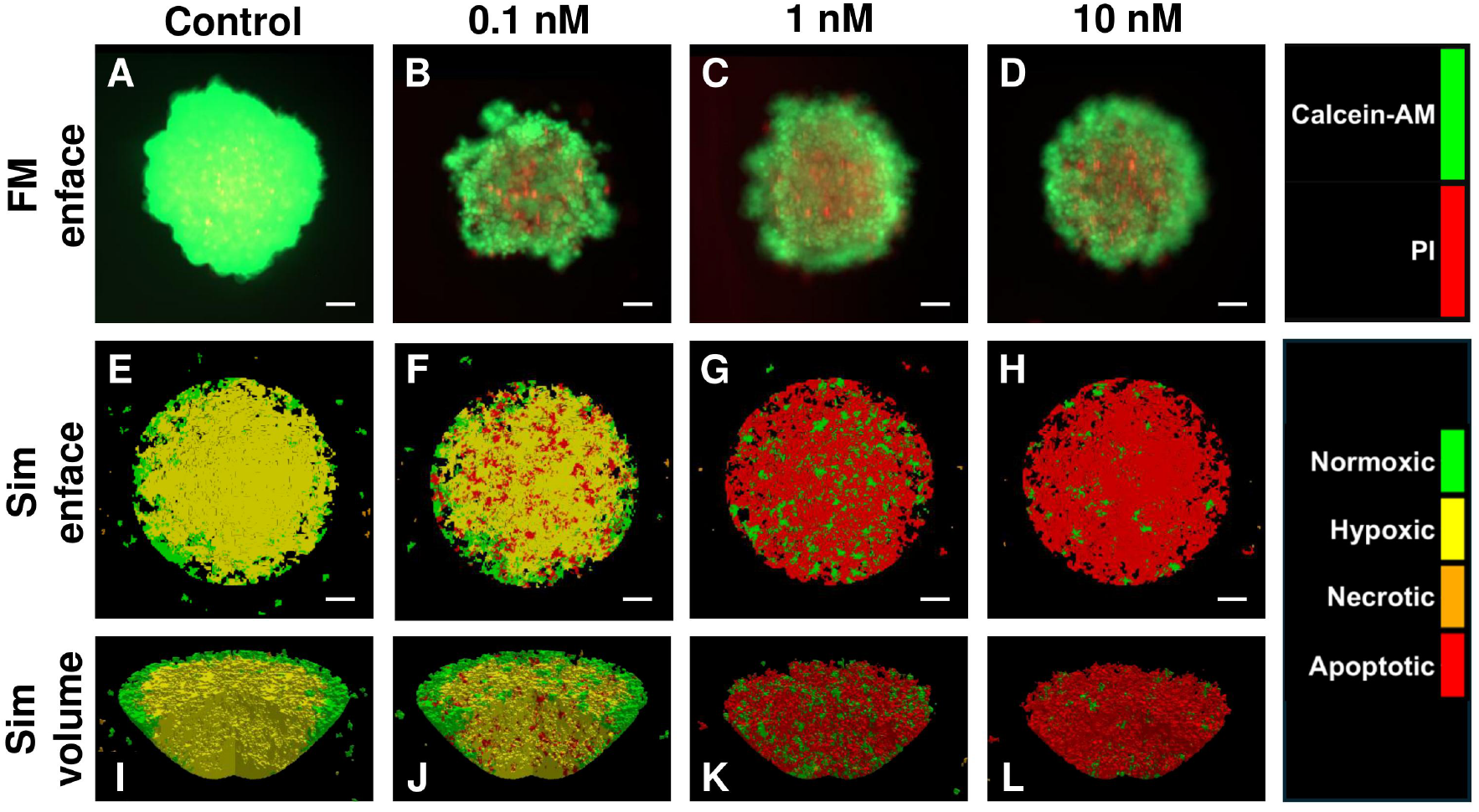
FM enface (first row), simulated enface (second row), and simulated volumetric (third row) images of spheroids treated with 0, 0.1, 1, and 10 nM DTX. A central slice has been cut out of the volumes. Colormaps for FM (red: PI, green: calcein-AM); and simulation (red: *Apoptotic*, orange: *Necrotic*, yellow: *Hypoxic*, green: *Normoxic*). DTX: docetaxel; FM: fluorescence microscopy; PI: propidium iodide; calcein-AM: calcein acetoxymethyl; Sim: simulation. Scale bars: 100 *µ*m.

The longitudinal observation of post-treatment recovery in spheroid and monolayer cultures are shown in Figures 4 and 5, respectively. AB measurement was performed pre-treatment on day 1, immediately post-treatment on day 3, and throughout treatment recovery on days 5 and 8. In spheroid culture, normalized AB absorbance fluctuated between 87% and 132% of the pre-treatment absorbance (Fig. 4M), whereas corresponding measurements in monolayer culture had much larger variations between 49% and 1225% (Fig. 5M). Normalized AB absorbance in control spheroids and those treated with lower DTX doses of 0.1 nM and 1 nM DTX were largely comparable, with each dose achieving the highest measurement on a different day and colony formation appearing largely similar (Fig. 4). In contrast, normalized AB absorbance in spheroids treated with 10 nM DTX was slightly lower than all other doses but remained higher than pre-treatment levels at each measurement time (Fig. 4M). Clonogenic plating of cells from spheroids treated with 10 nM DTX revealed colonies that were recognizable but smaller than all other doses each day (Fig. 4D,H,L). In monolayer culture, control and 1 nM DTX treated cells had comparable normalized AB absorbance, but on days 3 and 5 cells treated with 0.1 nM DTX had much larger values before falling to a value between control and 1 nM DTX by day 8 (Fig. 5M). Nonetheless, colony formation by 0.1 nM DTX treated monolayer cells was significantly faster and larger than all other conditions (Fig. 5B,F,J). AB absorbance measured in monolayer cells treated with 10 nM DTX decreased from pre-treatment AB absorbance levels throughout the measurement period (Fig. 5M) and barely formed colonies by day 14 (Fig. 5L).

**Figure 4.**
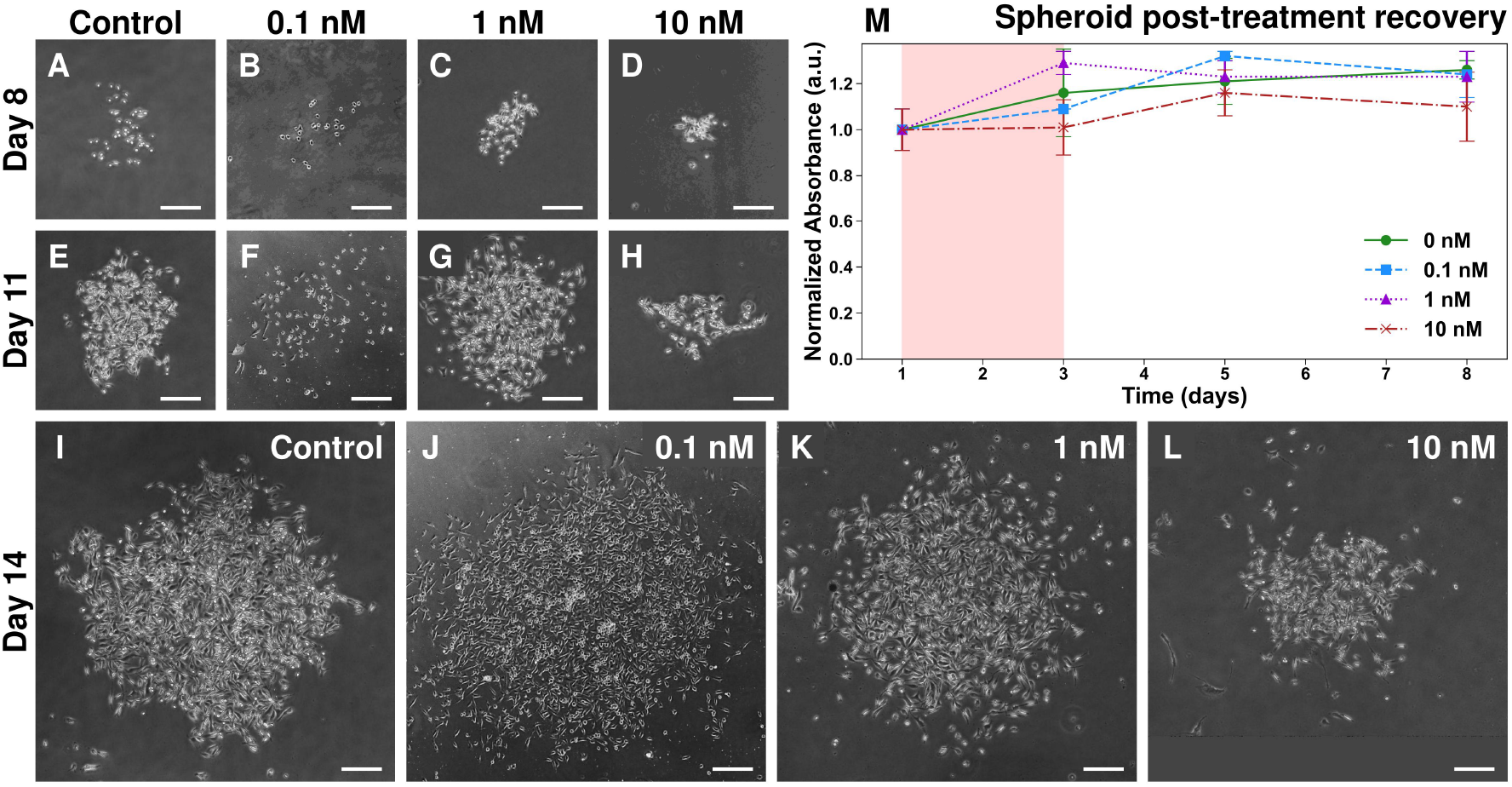
Longitudinal observation of post-treatment recovery in cells treated in spheroid culture with 0, 0.1, 1, and 10 nM DTX. Colony formation on day 8 (first row), day 11 (second row), and day 14 (third row) following removal of DTX and plating at low density in 2D on day 3. Normalized AB absorbance (top right) of intact spheroids treated with 0 nM (green circle, solid line), 0.1 nM (blue square, dashed line), 1 nM (purple triangle, dotted line), and 10 nM (orange cross, dot-dashed line) DTX. Exposure to DTX treatment occurred throughout pink shaded region. DTX: docetaxel; AB: Alamar Blue. Scale bars: 100 *µ*m.

**Figure 5.**
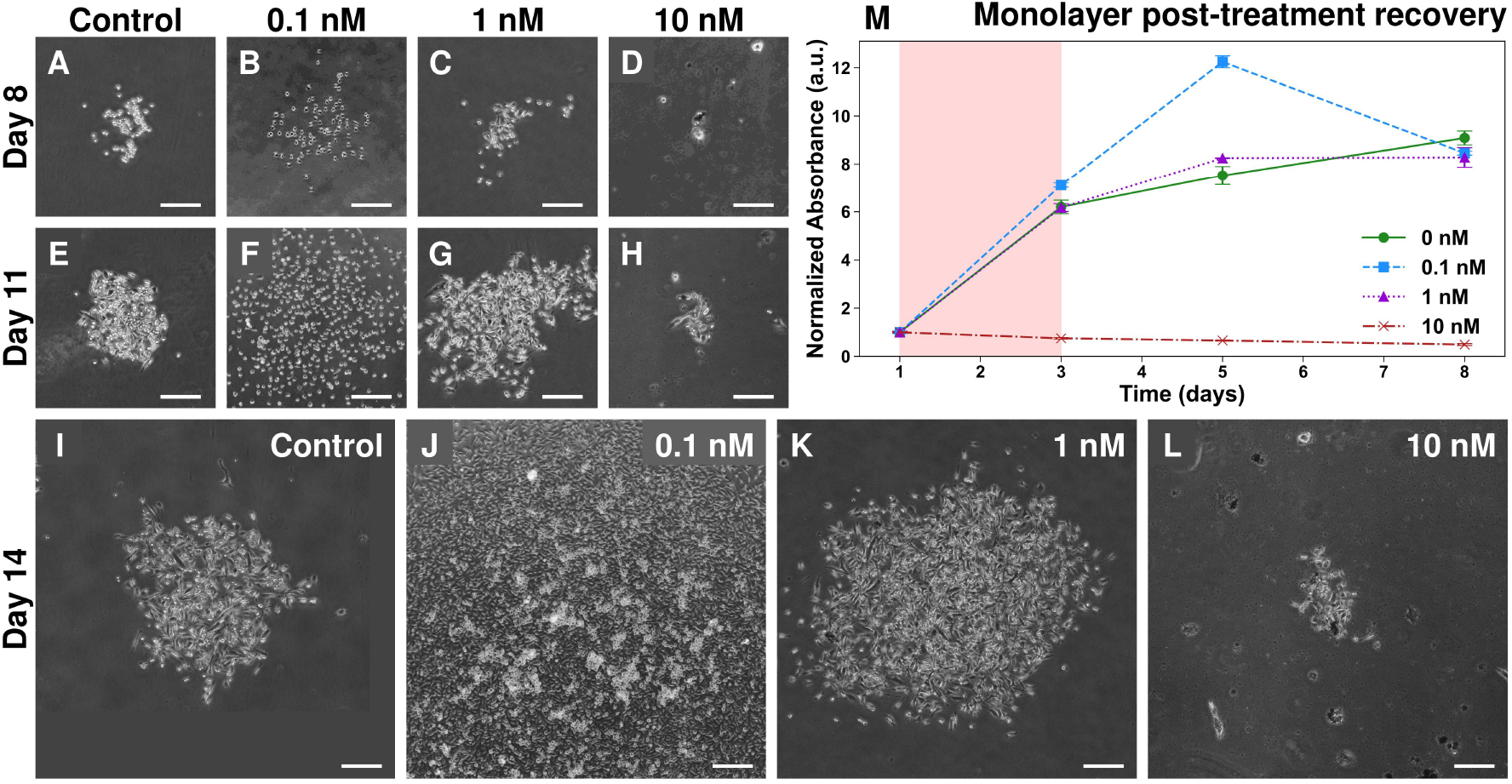
Longitudinal observation of post-treatment recovery in cells treated in monolayer culture with 0, 0.1, 1, and 10 nM DTX. Colony formation on day 8 (first row), day 11 (second row), and day 14 (third row) following removal of DTX and plating at low density in 2D on day 3. Normalized AB absorbance (top right) of monolayer culture treated with 0 nM (green circle, solid line), 0.1 nM (blue square, dashed line), 1 nM (purple triangle, dotted line), and 10 nM (orange cross, dot-dashed line) DTX. Exposure to DTX treatment occurred throughout pink shaded region. DTX: docetaxel; AB: Alamar Blue. Scale bars: 100 *µ*m.

## 4 Discussion

Clinically, prostate cancer patients are prescribed DTX according to body surface area, typically at 75 mg/m^2^ delivered by 1 hour of IV every three weeks [58]. Pharmacokinetic and pharmacodynamic study of this DTX regime observed an average peak plasma concentration of 4 *µ*M (3.26 *µ*g/mL) with an average 2.41 hour exposure to doses above 200 nM (0.16 *µ*g/mL) [59]. However, this dynamic delivery is difficult to reproduce with *in vitro* experiments, which generally expose cells to a constant DTX concentration for some period of time. Moreover, the dose limiting factor in DTX treatment is primarily hematologic toxicity [6]; as such, *in vitro* experiment cannot inform clinical delivery or dosage, but rather may be used to characterize and investigate the cellular response. Previous *in vitro* investigations treating prostate cancer cells with DTX have employed a wide variety of exposure times between 2 hours [60] and three weeks [61], with DTX doses ranging seven orders of magnitude from 6.4 pM [62] to 96 *µ*M [63]. Here, we exposed PC3 cells cultured in monolayers and spheroids to 0.1, 1, and 10 nM DTX for 48 hours. We assessed end-of-treatment survival and post-treatment recovery with clonogenic assay and AB proliferation assay.

### 4.1 End-of-treatment survival

As a disease, cancer is defined by uncontrolled cellular proliferation [64]. Since clonogenic assay directly and uniquely measures the ability of treated cells to continually divide successfully, it is particularly well suited for evaluating the success of cancer treatment. Clonogenic assay was first used to measure radiation effect in the field of radiobiology, where it remains the gold standard [65]. That said, its use has broadened to measure the effect of chemotherapies and other drug treatments [49]. Nonetheless, there has been limited clonogenic investigation of DTX-treated monolayers of prostate cancer cells, and to the best of our knowledge, none with spheroids.

We observed a higher clonogenic EC_50_ of 0.76 nM DTX in PC3 monolayer culture compared to 0.45 nM DTX in spheroid culture. However, cells cultured in monolayers were more susceptible to high doses of DTX and demonstrated approximately 2% clonogenic survival at 10 nM (Fig. 2A). Counterintuitively, after a longer exposure of three or four days compared to our 48 hours, Lange et al. [43] found a larger clonogenic EC_50_ of 1 nM DTX in PC3 cells cultured in monolayer. Likewise, clonogenic survival of PC3 monolayer culture treated with DTX for longer durations has yielded larger SF values of 93% at 0.1 nM after 10 days [44] and 57% at 1 nM after 72 hours [45], compared to our observations of 83% and 42% at 0.1 nM and 1 nM, respectively (Fig. 2A). However, by treating suspension culture of PC3 cells with DTX 2 hours, Karshafian et al. [60] and Almasri and Karshafian [66] observed similar clonogenic SF values at 0.1 nM of 81% and 75%, respectively. Linke et al. [67] observed equivalently low clonogenic survival of PC3 monolayers at 10 nM, albeit with 24 hours of post-treatment recovery before clonogenic plating. As with variation in measured survival within PC3 cell culture experiments, survival has also been found to vary by prostate cancer cell line [43, 45]. In particular, clonogenic assay has also been performed with DTX-treated monolayer cultures of LNCaP [43, 45, 68], LNCaP-derived C4-2 [69] and C4-2B [61], DU145 [43, 45, 67, 70, 71], VCaP [68], RWPE-2 [72], and 22Rv1 [73] prostate cancer cell lines. Notably, comparison between cell lines may be complicated as investigation of breast cancer cell lines has revealed cell-line-specific variation in the mechanism of DTX-induced cell death [74], potentially due to differences in regulation of proliferation, contact inhibition, or cell cycle checkpoints.

For comparison with clonogenic assay, one of our AB measurement times was at end-of-treatment. Similar investigations of PC3 monolayers treated for 48 hours with DTX have observed a wide range of EC_50_ values using various proliferation assays, including 0.984 nM [75] and approximately 10 nM [76] via MTT, and 33.89 nM via CCK-8 [77]. Monteverde et al. [75] identified a minimum survival of 10% at 2 nM DTX by MTT relative to control, whereas we found end-of-treatment normalized AB absorbance in monolayers was 12% at 10 nM and comparable to control at 1 nM (Fig. 5 M). In comparison, end-of-treatment AB measurement fluctuated much less in spheroid culture and spheroids treated with 10 nM DTX demonstrated 87% absorbance compared to control. Pereira et al. [78] similarly evaluated PC3 spheroids treated with DTX for 48 hours by AB and observed approximately 55% absorbance at 1 nM and 10 nM compared to control. Other investigators have similarly performed proliferation assay and observed higher DTX survival in spheroids compared to monolayers with LNCaP [79, 80], DU145 [63, 80, 81], and PC3 [80] cell lines.

Investigations of DTX-treated prostate cancer cells via proliferation assay far outnumber those via clonogenic assay. We suspect one reason for the disparity is that, even for investigation of monolayer culture, clonogenic assay is an arduous method that demands trial and error. Moreover, we could not find a single investigation that performed clonogenic assay of DTX-treated prostate cancer spheroids, likely because the nature of 3D cell culture makes clonogenic assay much harder to perform. While AB measurement is performed with intact spheroids, longitudinal observation of colony formation by clonogenic assay requires spheroid disaggregation by a physically stressful process involving aggressive enzymatic dissociation by trypsin, mechanical vortexing, and centrifuging.

Furthermore, it is experimentally challenging to employ clonogenic and proliferation assays with the same doses of DTX. Experiments employing clonogenic assay have exposed PC3 monolayers to DTX concentrations up to 0.1 nM [66], 0.5 nM [44], 5 nM [43], 9 nM [60], and 10 nM [45]; in comparison, proliferation assays performed with PC3 monolayers used much larger maximal DTX concentrations of 10 nM [76], 62.5 nM [77], 100 nM [75], 250 nM [82], and 1 *µ*M [67]. Proliferation assays of PC3 spheroids have employed even higher concentrations of 200 nM [80], 2.5 *µ*M [82], and 10 *µ*M [78]. The discrepancy in DTX dosage required to differentiate the effect of treatment via clonogenic and proliferation assays may be due to treated cells that lack reproductive viability while remaining metabolically active, for example by senescence. Moreover, longitudinal observation via clonogenic assay isolates viability measurement to the initial population of cells. In contrast, longitudinal proliferation assay measurements are complicated by detecting the cellular activity of both the initially surviving cells as well as their progeny.

For example, Zemskova et al. [72] performed clonogenic assay with RWPE-2 monolayers with doses up to 20 nM DTX, but all proliferation assays were performed via MTT with 100 nM DTX. Fujiike et al. [63] attempted to investigate DTX treatment of monolayer and spheroid DU145 culture with both AB and clonogenic assay. By AB proliferation assay, they found that EC_50_ values were approximately 10-fold higher in spheroids compared to monolayer culture. However, they observed no colony formation via clonogenic assay by plating 300 viable cells in 6-well plates after 24 hours of treatment with doses ranging from 1 to 96 *µ*M DTX. In contrast, to perform clonogenic assay for 10 nM monolayer-treated and spheroid-treated cells, we plated 2,400 cells and 12,000 cells in 6-well plates, respectively. Despite the higher SF of spheroid culture at 10 nM, the process of disaggregation lowers spheroid PE such that a higher number of cells must be plated to produce a statistically sound number of colonies. This restricts the upper dose limit accessible to analysis by clonogenic assay, especially for spheroid culture. Potentially, our AB analysis was limited by imposing maximum doses that were also feasible for clonogenic assay, resulting in little variation with dose in spheroids at end-of-treatment (Fig. 2B). Nonetheless, long exposure time to higher doses is not clinically achievable [59]; rather, here we are interested in the recovery and clonogenic potential of prostate cancer cells following treatment.

### 4.2 Post-treatment recovery

In addition to pre-treatment and end-of-treatment measurements on days 1 and 3, we also measured AB absorbance of monolayer (Fig. 5M) and spheroid (Fig. 4M) cultures in their treatment vessels throughout post-treatment recovery on days 5 and 8. After plating clonogenic assay at end-of-treatment, we documented longitudinal colony formation for qualitative comparison with AB trends (Fig. 4, 5). Some post-treatment measurements have been made previously in DTX-treated PC3 monolayers at 24 hours [67] and 48 hours [82, 83] post-treatment, and with C4-2B cells after 72 hours [61]. However, these post-treatment measurements were not designed to investigate recovery, but rather are a byproduct of sensitization agent delivery [83] or observing drug synergism [61].

Bromma et al. [62] performed longitudinal post-treatment observation of 24 hour DTX-treated LNCaP monolayer and spheroids using CellTiter-Glo 3D proliferation assay at 2 days post-treatment and at 6 and 13 days post-treatment, respectively. They estimated spheroid volume regularly from enface bright-field microscopy images, but they only investigated one dose of DTX and did not measure proliferation assay at pre-treatment or end-of-treatment.

We observed better post-treatment recovery in spheroids, with little fluctuation in AB measurement by DTX dose (Fig. 4M) and higher clonogenic survival at higher doses (Fig. 2A). In contrast, 10 nM treated monolayer culture never recovered to pre-treatment AB measurement (Fig. 5M). Not only was monolayer culture more susceptible to high doses, but a low dose of 0.1 nM appeared to stimulate rather than inhibit monolayer growth (Fig. 5M). We observed a similar effect to a lesser extent in spheroid culture (Fig. 4M). Colony formation occurred faster in 0.1 nM treated monolayer cells than any other condition, followed by 0.1 nM treated spheroids (Fig. 4, 5). Despite demonstrating an increased speed of colony formation, monolayer and spheroid culture treated with 0.1 nM DTX produced fewer colonies than control (Fig. 2). Compared to untreated monolayers, other investigations of prostate cancer cells have observed higher proliferation after treatment for 48 hours with doses 0.1 nM DTX and lower in PC3 monolayers [75, 76, 82] and 24 hours with doses 1 nM DTX and lower in RWPE-2 monolayers [72].

These observations are consistent with a controversial phenomenon called hormesis [84], whereby an agent exhibits a biphasic dose response with growth inhibition at high doses and growth stimulation at low doses. The concept of hormesis was introduced in the 1880s by Schulz [85] and coined in 1943 by Southam and Ehrlich [86]. It has since been extensively documented and debated [87, 88], particularly in the area of radiobiology [89, 90]. However, chemotherapeutic hormesis has not been well characterized, especially outside of immortalized cell lines [91]. In post hoc analysis of their previous work [92], Yoshimasu et al. [93] identified negative inhibition rates of DTX with *in vitro* histoculture drug response assay (HDRA) of samples from patients with non-small-cell lung cancer (NSCLC). They suggested hormesis as the cause, but employed a large dose of 124 *µ*M (100 *µ*g/mL) DTX for a duration of 7 days. They analyzed the same specimens with a range of paclitaxel doses and observed growth stimulation at low doses [94], but had not designed the experiment around the low dose range required to validate hormetic behavior [93]. Chang et al. [91] observed paclitaxel-induced hormesis by MTS in NOS3 ovarian cancer cell monolayers, but did not perform clonogenic assay for comparison. In general, hormesis has been widely demonstrated in cell culture [87], but investigations have been critiqued for lacking appropriate experimental controls or sufficient sample of doses [88]. Our study was also not designed to reveal DTX hormesis, but may suggest the first measurement of its kind in prostate cancer spheroid culture, which motivates further investigation.

### 4.3 Simulation

At end-of-treatment, simulated clonogenic and AB values were similar to experiment (Fig. 2B). Notably, simulated AB also demonstrated hormesis with higher simulated proliferation observed at 0.1 nM DTX compared to control; however, simulated and experimental clonogenic results agreed well and decreased monotonically with dose (Fig. 2B). In this way, simulated spheroids visually indicated cells that are destined for reproductive death, whereas FM images reveal transient cellular health that rather visualizes AB proliferation (Fig. 3). As previously suggested [50], cellular activity may be increased following treatment due to the numerous signaling pathways activated by cell repair [95] and death [56, 96]. Simulated AB values were calculated under the assumption that following an inactive period of cell cycle arrest, successfully treated cells undergo an active apoptotic process. However, these cells have lost clonogenic potential and thus do not contribute to simulated clonogenic survival. Removing transient apoptotic activity from simulated AB calculations to represent growth alone removes the hormetic effect and calls into the question the mechanism of higher measured AB proliferation under low DTX dose (Fig. 2B). The low dose growth stimulation measured by proliferation assay may be a byproduct of cellular activity associated with DTX-induced cell death or ineffectual repair.

Using a PDE model, Hinow et al. [97] explored drug-specific parameters and similarly observed increased proliferation in a simulated *in vivo* tumor following treatment with a DTX-like drug. However, the growth stimulation was caused by an imbalance of drug-induced cell killing and drug decay rather than a hormetic effect of low dose. DTX-specific simulations have been performed with an ABM of monolayer culture [98], an optimal control framework [99], and a PDE model [54]. Kaura et al. [100] investigated paclitaxel treatment with a 2D CPM based on the spheroid model developed previously by the authors [33]. They observed FM [33] and simulated [33, 100] images with increased paclitaxel effect along the spheroid periphery, in contrast to the cell death observed centrally in our FM and throughout our simulated images (Fig. 3). Similar to the wide range of conflicting experimental *in vitro* DTX measurements, ABM simulations of chemotherapy treatment have made contradictory conclusions that periodic drug administration [101] or constant dosing [102] is more effective.

### 4.4 Limitations

To the best of our knowledge, there has not been previous clonogenic measurement of DTX-treated spheroid culture or longitudinal observation of post-treatment DTX-dose dependent effects in prostate cancer monolayer or spheroid culture. Still, the time scale of our observation was limited by monolayer culture becoming confluent by day 8 of the experiment. To ensure a sufficient number of cells at each AB measurement, we seeded monolayer culture at different cell densities by dose (Table 1), and normalized each dose to its own pre-treatment measurement. Unlike many other investigations that diluted DTX in cytotoxic DMSO, we employed 0.9% saline for our dilutions, but as with the other studies, we present normalized results that we expect to remove any effect of dilution medium. While the upper dose limit was restricted by clonogenic plating of spheroid culture, AB measurement limited the number of doses investigated due to the laborious process of removing DTX from spheroid culture gels by four serial dilutions. For each dilution of each gel, 5 mL of RPMI was added dropwise to avoid mechanically agitating the spheroids. Nevertheless, cell loss was unavoidable, but increased post-treatment AB absorbance from pre-treatment levels indicated spheroids continued to grow (Fig. 4M). However, interpretation of longitudinal AB absorbance in spheroids may be complicated by our previous observation of cyclical AB measurements [53]. Clonogenic plating of spheroid culture is also limited by removing the cells from the spheroid structure, which changes their phenotype [20, 22] and potentially their post-treatment response to DTX.

In contrast to our *in silico* model, previous models of chemotherapy treatment have simulated cell cycle phases and arrest [103], repair [33], and resistance [101]. Nonetheless, our model incorporates clonogenic survival measurements made in spheroid culture that integrate these complicated biological phenomena into a binary probability of reproductive survival or death. A direction for future work involving the model could include the evaluation of drug synergy and combination therapies, as has been previously simulated with DTX and other chemotherapeutics [33, 100], anti-angiogenic agent [54], ADT [99], targeted therapy [104], and immunotherapy [105]. Combination therapy with radiation is of particular interest because DTX arrests and synchronizes cells in G2-M [14], which is the most radiosensitive phase in the cell cycle [106].

## 5 Conclusion

In conclusion, we assessed end-of-treatment survival and post-treatment recovery via proliferation and clonogenic assays in PC3 cells cultured in monolayers and spheroids that were exposed to 0.1, 1, and 10 nM DTX for 48 hours. To the best of our knowledge, we performed the first clonogenic measurement of DTX-treated prostate tumor spheroids and dose-dependent longitudinal observation of post-treatment recovery. We observed growth stimulation by low dose DTX in monolayer and, to a lesser extent, spheroid culture via AB proliferation assay. While clonogenic assay did not measure improved survival under low dose treatment, observation of colony formation demonstrated faster and larger colony formation by surviving cells. With parameters informed by experimental clonogenic assay data, simulation of a novel 3D ABM of DTX treatment suggested the hormetic effect measured by proliferation assay may have been influenced by the active process of apoptosis. Future work exploring DTX hormesis should sample a sufficient number of doses with careful attention to experimental controls.

## Acknowledgments

The authors would like to gratefully acknowledge Dr. Mohammad Kohandel for generous use of his laboratory equipment. We thank Dr. Brian Ingalls and Atiyeh Ahmadi for their expertise and use of their fluorescent microscope, and Dr. Qing-Bin Lu and Olya Changizi for access and assistance with their microplate reader.

## Data availability

Data underlying the results presented in this paper are not publicly available at this time but may be obtained from the authors upon reasonable request.

## Code availability

The model code used for simulation is publicly available on GitHub at github.com/skswanso/CPM.git.

